# Precise identification of cell states altered in disease with healthy single-cell references

**DOI:** 10.1101/2022.11.10.515939

**Authors:** Emma Dann, Sarah A. Teichmann, John C. Marioni

## Abstract

Single cell genomics is a powerful tool to distinguish altered cell states in disease tissue samples, through joint analysis with healthy reference datasets. Collections of data from healthy individuals are being integrated in cell atlases that provide a comprehensive view of cellular phenotypes in a tissue. However, it remains unclear whether atlas datasets are suitable references for disease-state identification, or whether matched control samples should be employed, to minimise false discoveries driven by biological and technical confounders. Here we quantitatively compare the use of atlas and control datasets as references for identification of disease-associated cell states, on simulations and real disease scRNA-seq datasets. We find that reliance on a single type of reference dataset introduces false positives. Conversely, using an atlas dataset as reference for latent space learning followed by differential analysis against a matched control dataset leads to precise identification of disease-associated cell states. We show that, when an atlas dataset is available, it is possible to reduce the number of control samples without increasing the rate of false discoveries. Using a cell atlas of blood cells from 12 studies to contextualise data from a case-control COVID-19 cohort, we sensitively detect cell states associated with infection, and distinguish heterogeneous pathological cell states associated with distinct clinical severities. Our analysis provides guiding principles for design of disease cohort studies and efficient use of cell atlases within the Human Cell Atlas.

## Introduction

High-dimensional tissue profiling of healthy and disease samples with single-cell genomics enables the characterization of cellular phenotypes that are altered in disease (Lindeboom, Regev and Teichmann, 2021). Precise identification of such phenotypes can yield new mechanistic insights into pathogenesis, novel biomarkers and potential drug targets (Reyfman *et al*., 2019; Velmeshev *et al*., 2019; Adams *et al*., 2020; Elmentaite *et al*., 2020; Reichart *et al*., 2022; Richard K. Perez *et al*., 2022), with drugs against targets first identified using single-cell analysis starting to enter clinical trials (Eisenstein, 2022).

To identify altered cell states, joint analysis of single-cell RNA-sequencing (scRNA-seq) data of diseased tissues and a healthy reference is the standard practice. Typically, cellular profiles from different conditions are first integrated into a common phenotypic latent space, matching common cell types and minimising technical differences (Hao *et al*., 2021; Lotfollahi *et al*., 2022). Then, healthy and diseased cells in matched cell states are contrasted in differential analysis, to identify differences in gene expression patterns or in cellular composition (Burkhardt *et al*., 2021; Skinnider *et al*., 2021; Zhao *et al*., 2021; Dann *et al*., 2022). While the choice of method used for both steps can be impactful, appropriate selection of the healthy reference dataset is crucial for precise identification of true disease-associated states.

The profiling of collections of healthy tissue samples by the research community has enabled the generation of large, harmonised collections of data from multiple organs, leading to the rapid expansion of the Human Cell Atlas (http://data.humancellatlas.org/). Indeed, for certain tissues (e.g., lung and blood) there now exist millions of cells profiled using a variety of technologies from hundreds to thousands of individuals. Computational analyses allow these datasets to be meaningfully integrated, thus providing a comprehensive view of cell phenotypes in a tissue, while effectively minimising variation driven by experimental protocols. Nevertheless, the biological and technical characteristics of the samples included in an atlas might differ greatly from those of the disease cohort of interest. This could lead to false discoveries in differential analysis, if confounding factors are unknown or not appropriately handled in statistical testing.

Instead of using externally-generated atlases as a reference against which to compare disease samples, a matched control dataset could be used, where healthy tissue samples are selected to ‘match’ the disease samples in terms of cohort size, demographics, sample collection and processing protocols (ideally with healthy and diseased samples processed in parallel). This choice of reference minimises the risk of false positives driven by confounding effects. However, collection of a large number of healthy control samples is not always practical or possible, especially in human studies. Moreover, using a relatively small number of samples for the integration step increases the risk of missing rare cell states, and over-interpreting sample-specific noise. Understanding how the features of the reference dataset affect the ability to identify disease-associated cell states will guide effective data re-use, design of disease studies, and future cell atlasing efforts.

Here we compare the use of atlas and control datasets as references for identification of disease-associated cell states, showing that reliance on a single type of reference dataset introduces false positives. In contrast, combined use of an atlas dataset as reference for latent embedding and of a control dataset as reference for differential analysis leads to precise and sensitive identification of putative disease cell states, with important implications for both experimental design and utilisation of single-cell disease cohorts.

## Results

### Reference design for identification of disease-associated cell states

To optimize the selection of a reference dataset for identification of disease-associated cell states, we considered the attributes of the disease and reference datasets that are commonly jointly analysed in scRNA-seq studies (figure 1A). Throughout this study we use the term “cell state” to define a group of cells that are more transcriptionally similar to each other than to other cells in the same tissue - in practice a cell state might represent a neighbourhood of cells, the output of a clustering algorithm or cell type annotation. In a disease dataset, biological samples typically originate from tens of individuals from a relatively homogeneous population (e.g. recruited from the same hospital), the same experimental protocol is used across samples for dissociation, library prep and sequencing (or experiments are designed to minimise confounding with cohort-specific variables). We define a healthy reference dataset as a *control* if it matches the disease dataset in terms of cohort characteristics and experimental protocols. We define a reference dataset as an *atlas* if it aggregates data from hundreds to thousands of individuals from multiple cohorts, profiled with a variety of experimental protocols. Typically, such integrated datasets capture a larger variety of healthy cell states compared to smaller cohorts.

**Figure 1:**
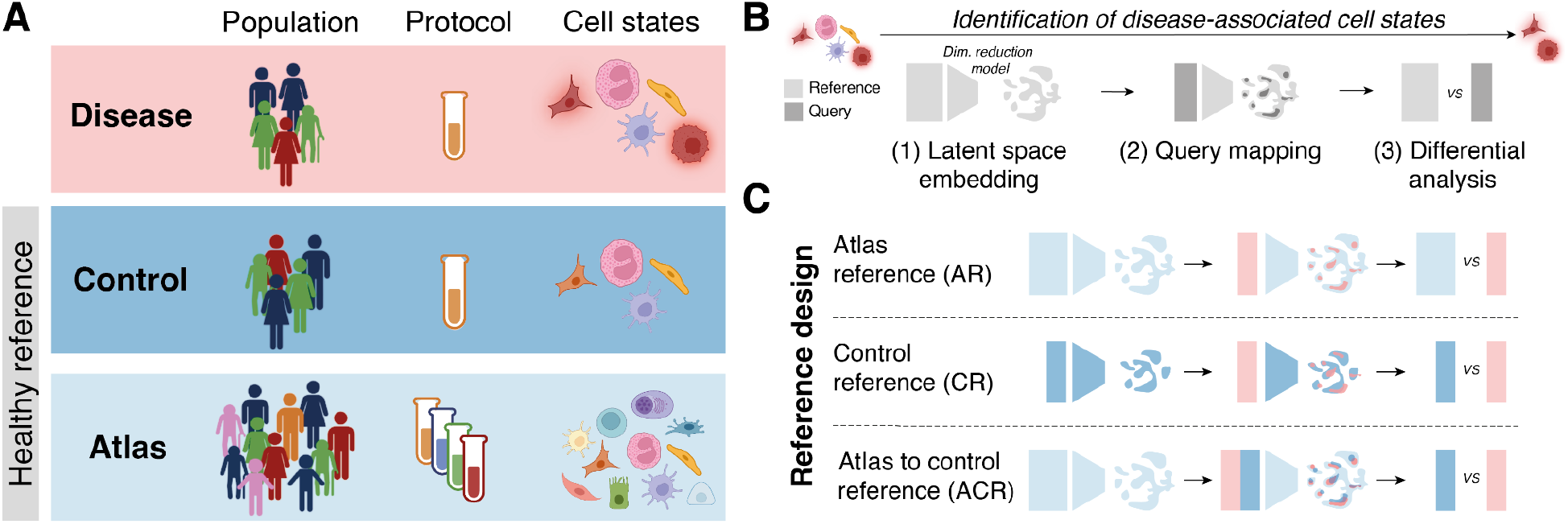
Using healthy reference datasets to discover disease-associated cell states. (**A**) Schematic of attributes of disease, control and atlas datasets, with regards to population-level variation, experimental protocols, and heterogeneity of cell states captured. (**B**) Schematic of analysis workflow to detect disease-associated cell states: a dimensionality reduction model is trained on a healthy reference dataset (step 1), then the query dataset, including the disease dataset, is mapped to the reference model with transfer learning (step 2), finally differential analysis is performed to contrast matched cell states from healthy and disease samples. **(C)** Schematic of reference design options tested in this study, following the workflow in (B): using the atlas dataset as reference (light blue), the control dataset as reference (dark blue) or both.

We consider the following workflow to identify disease-associated cell states (figure 1B). First, a dimensionality reduction model is trained on the dataset from healthy tissues (embedding reference dataset), to minimise batch effects while learning a latent space representative of cellular phenotypes in the tissue. Then, the model trained on the embedding reference is used to map the query dataset, including the disease samples, to the same latent space using transfer learning (TL) (Hao *et al*., 2021; Lotfollahi *et al*., 2022). Finally, differential analysis comparing cells between disease samples and healthy samples (differential analysis reference) is used to distinguish disease-associated states. With this workflow, we identify three alternatives for selection of a reference dataset (*reference design*) (figure 1C): i) the atlas reference design (AR design) or ii) the control reference design (CR design), where either is used as embedding reference and as differential analysis reference, and iii) a design where an atlas and a control dataset are used in different steps of the workflow (ACR design). In this analytical design, the atlas dataset is used as the embedding reference; subsequently, both the disease and the control datasets are mapped to the same latent space; finally, differential analysis is performed contrasting the disease dataset to the control dataset only.

In the following sections, we quantitatively assess the ability of these three designs (illustrated in Fig 1C) to identify disease-specific cell states in simulations and real data.

### Simulations show precise detection of out-of-reference cell states with combined use of atlas and control datasets

To compare reference designs in a scenario with ground-truth, we simulated the attributes of atlas, control and disease datasets by splitting scRNA-seq data from real studies (figure 2A). We collected publicly available data from 13 studies that profiled healthy peripheral blood mononuclear cells (PBMCs), comprising profiles from 1248 donors, which we harmonised to obtain a consistent cell type annotation (see Methods). We select cells from one study (29 donors) and randomly split the donors to simulate the disease dataset (16 donors) and the control dataset (13 donors). Using different donors from the same study ensures that cohort demographics and experimental protocols are matched between disease and control datasets, while donor and library effects present in real data are maintained. The cells from the 12 remaining studies (1219 donors) form the atlas dataset. To simulate the presence of a cell population specific to the disease dataset - which we define as an *out-of-reference* (OOR) state, we select an annotated cell type and remove cells with that label in the control and atlas dataset. Overall, we tested the ability to identify 15 different cell types as the OOR state.

**Figure 2:**
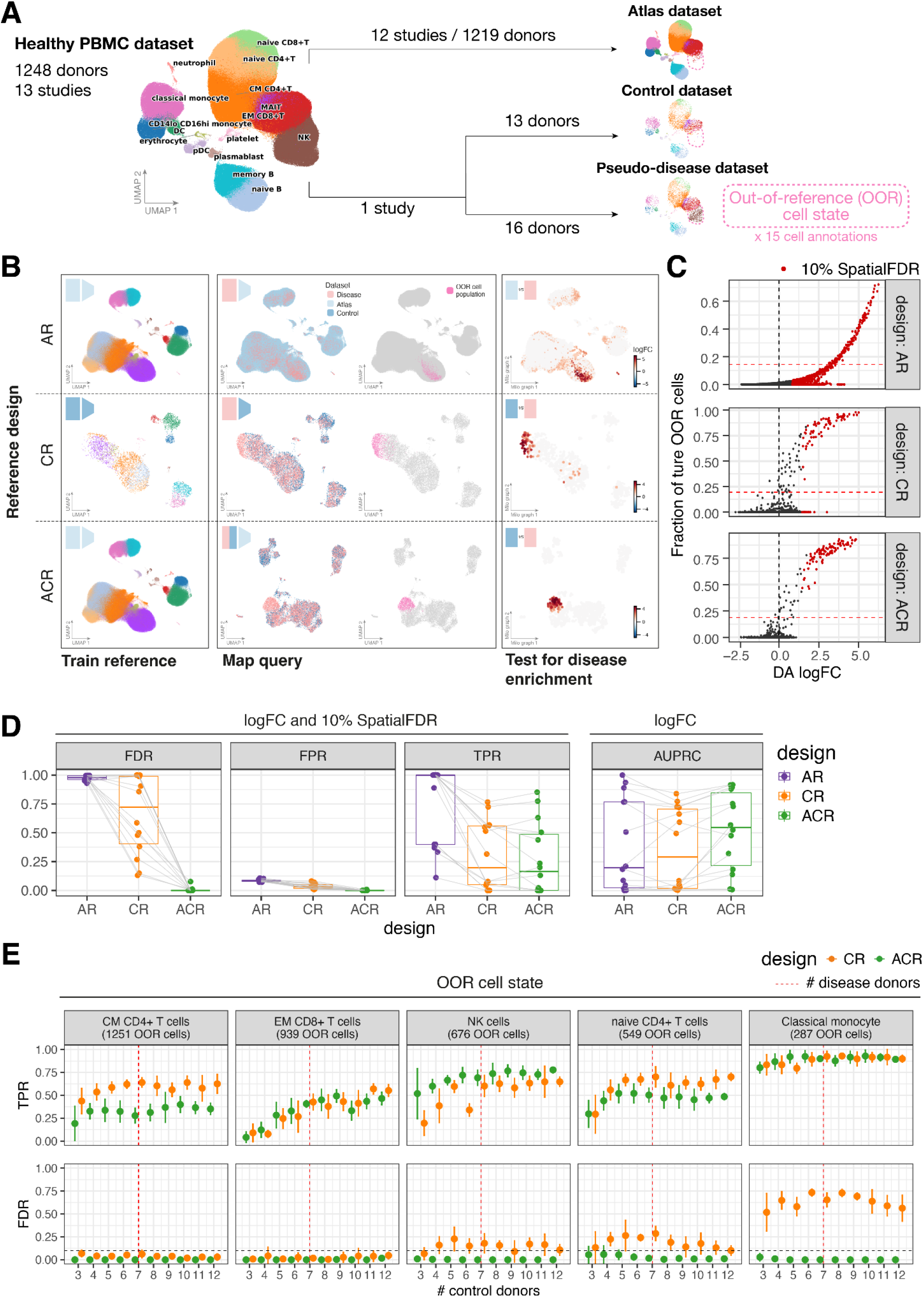
Combined use of atlas and control datasets as references for precise and sensitive detection of out-of-reference cell states. **(A)** Schematic of the strategy used to simulate ground-truth out-of-reference (OOR) cell states in real data from healthy human peripheral blood mononuclear cells (PBMCs), splitting in atlas dataset (513565 cells), control dataset (5671 cells) and pseudo-disease dataset (7505 cells). We tested simulations alternatively using 15 annotated cell states as OOR cell state. **(B)** Example outcome of query-to-reference mapping and differential analysis with different reference designs: (left) UMAP embedding of scVI latent space learnt on embedding reference dataset, points are colored by cell type clusters (as in A) and the icons in the top left corner indicate the type of embedding reference dataset used; (center) UMAP embedding of cells from differential analysis reference and disease datasets on scVI latent space learnt from embedding reference dataset, colored by type of dataset and to highlight (in pink) the OOR cell state; (right) Milo neighbourhood graph visualisation of differential abundance testing results: each point represents a neighbourhood, points are colored by the log-Fold Change (logFC) in cell abundance between disease and reference cells. Only neighbourhoods where significant enrichment in disease cells (10% SpatialFDR and log-Fold Change > 0) was detected are colored. Points are positioned based on the coordinates in UMAP embedding of the neighbourhood index cell, the size of points is proportional to the number of cells in the neighbourhood. **(C)** Scatterplot of differential abundance log-Fold Change against fraction of perturbation-specific cells for each neighbourhood for the simulation shown in C. Each plot represents a different reference design. Colored points indicate neighbourhoods where significant enrichment in disease cells (10% SpatialFDR and log-Fold Change > 0) was detected. **(D)** Quantitative comparison of performance of reference designs in detection of OOR cell states (neighbourhoods where the fraction of OOR cells is higher than 20% of the maximum fraction for that simulation). To compare performance considering the logFC and confidence (10% SpatialFDR), we measured the false discovery rate (FDR), false positive rate (FPR) and true positive rate (TPR). To compare performance using logFC only as a metric for prioritization, we measure the Area Under the Precision-Recall Curve (AUPRC). Points represent simulations with different OOR states. Tests on the same simulated data are connected. **(E)** Robustness to size of control cohort with ACR and CR designs: mean true positive rate (TPR) and false discovery rate (FDR) for simulations with increasing number of donors in the control dataset (x-axis), using atlas to control reference design (ACR, green) or control reference design (CR, orange). The error bar shows the standard deviation over five simulations using a different sample of donors. In these simulations 7 donors were used in the disease dataset. Results from simulations with five different OOR cell states are shown, selected by top mean TPR across designs in (D).

We apply the following workflow to identify the OOR state: we learn a latent space embedding on the reference of choice (atlas or control) using the scVI model (Lopez *et al*., 2018) (figure 2B, left). Then, we use transfer learning with scArches (Lotfollahi *et al*., 2022) to map the query dataset(s) to the trained scVI model. This places cells from the disease and reference datasets in the same latent space, which captures common cell states (figure 2B, centre). Of note, with the ACR design, the atlas dataset is used only to train the latent embedding model, but after mapping with scArches only disease and control datasets are used for differential analysis. This reduces the computational burden of handling a dataset of hundreds of thousands of cells. Finally, we use neighbourhood-level differential abundance (DA) testing with Milo (Dann *et al*., 2022) to identify cell states enriched in cells from the disease dataset (Figure 2B, right), where neighbourhoods with SpatialFDR < 10% and a log-Fold Change (logFC) in abundance > 0 are defined as predicted OOR cell states.

Across simulations with different OOR states, we found that combining the use of the atlas and control dataset (ACR design) led to sensitive detection of neighbourhoods with a high fraction of OOR cells (Figure 2C-D, suppl. figure 1). Conversely the AR design led to an inflated number of false positives, where significant enrichment was also detected when the fraction of unseen cells is low or 0. Using only the control dataset led to more balanced logFCs, but still a higher false positive rate compared to the ACR design. Only the ACR design maintained FDR control across simulations (i.e. it rarely identifies enriched neighbourhoods with a small fraction of unseen cells) (Figure 2D). The ACR design also performs better when using just the logFC as a metric to prioritize neighbourhoods containing OOR cells, regardless of the p-value, by quantifying the area under the precision-recall curve (AUPRC) (Figure 2D, right). The overall variance in sensitivity across different simulations is mostly explained by the number of cells in the OOR state (suppl. figure 2). Notably, we found no significant difference in the quality of integration with different designs, in terms of conservation of cell types and batch removal (suppl. figure 3).

**Figure 3:**
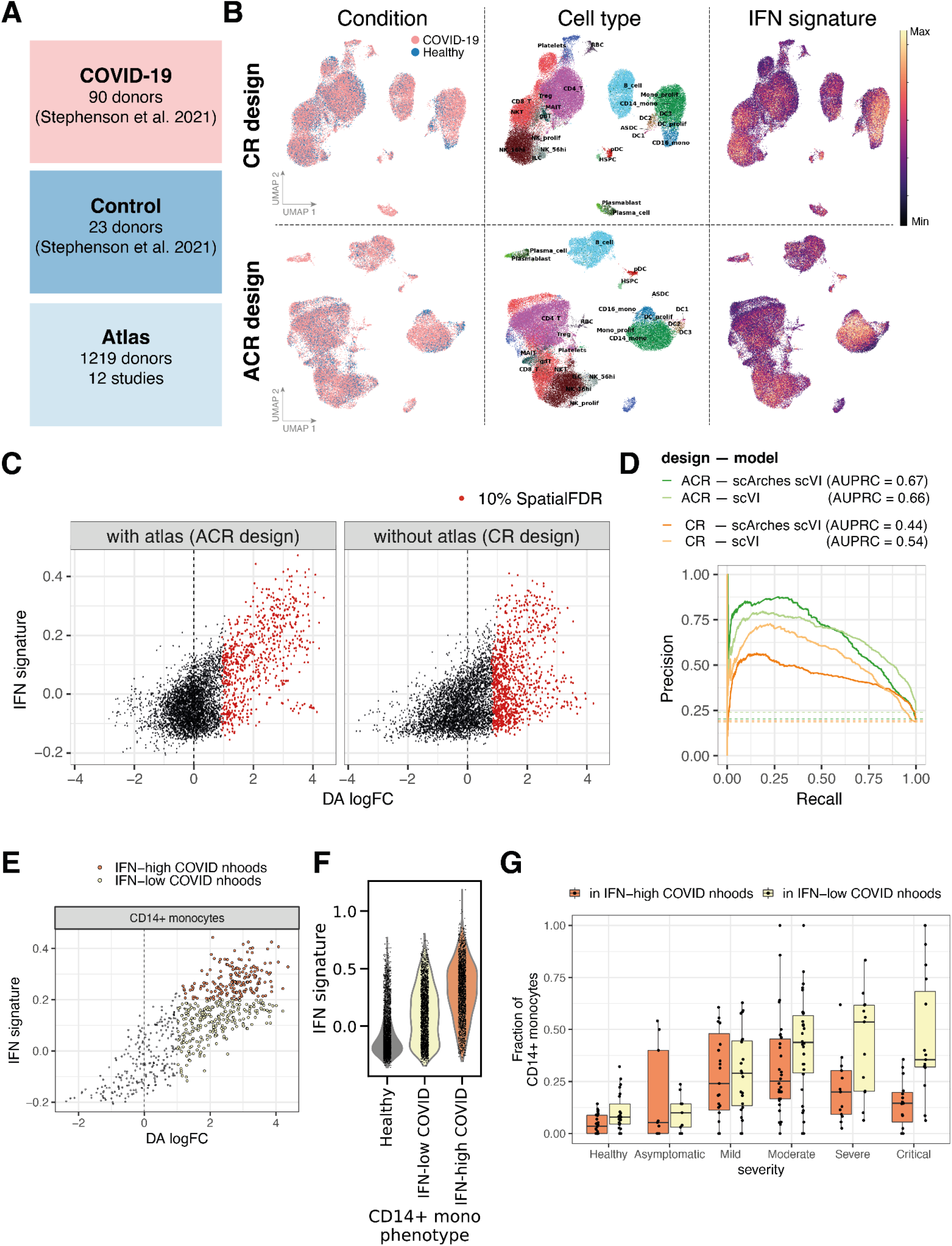
Detection of cell states associated with COVID-19 in a case-control cohort with a healthy atlas. **(A)** Overview of composition of disease, control and atlas dataset. **(B)** UMAP embedding of cells from COVID-19 and healthy datasets integrated with CR design (top) or ACR design (bottom). Cells are colored by disease condition (left), broad annotated cell type (middle) and expression of interferon (IFN) signature (right). **(C)** Scatterplot of neighbourhood differential abundance log-Fold Change (DA logFC) against the mean expression of IFN signature with ACR design (left) and CR design (right). Colored points indicate neighbourhoods where the enrichment in COVID-19 cells was significant (logFC > 0 and 10% SpatialFDR). **(D)** Precision-recall curves for detection of IFN-activated neighbourhoods with DA logFC for alternative designs (ACR or CR) and using de novo integration of reference and disease datasets (scVI) or transfer learning (scArches scVI). The area under the PR curve (AUPRC) is reported in the legend. The dotted lines denote the baseline value for the AUPRC, indicating the case of a random classifier. **(E)** Scatterplot of neighbourhood differential abundance log-Fold Change (DA logFC) against the mean expression of IFN signature with ACR design for neighbourhoods of CD14+ monocyte cells. Colored points indicate neighbourhoods where the enrichment in COVID-19 cells was significant (10% SpatialFDR). Neighbourhoods are colored by IFN phenotype. **(F)** Distribution of IFN signature score for cells belonging to neighbourhoods assigned to 3 alternative CD14+ phenotypes. **(G)** Distribution of COVID-19 enriched CD14+ phenotypes across patients with varying disease severity: each point represents a donor, the y-axis shows the fraction of all CD14+ monocytes in that donor showing IFN-high COVID-19 enriched phenotype (orange), and IFN-low COVID-19 enriched phenotype (yellow). The remaining fraction are monocytes with healthy phenotype (not shown).

Studies that use atlas datasets to identify disease-associated states (Hao *et al*., 2021; Lotfollahi *et al*., 2022, Sikkema et al. 2022) often omit differential testing and define OOR cells using two criteria. Firstly, OOR cells are expected to segregate from atlas cells in the latent space. In our simulations the OOR cell state is not consistently distinguished by clustering or by distance to reference cells in the latent space (figure 2B, suppl. figure 4). Secondly, different cell-level metrics for query-to-reference mapping quality are used to flag OOR cells, such as reconstruction error or uncertainty over a cell type classifier trained on the atlas. The assumption is that, if the disease-specific cell state was not seen during training of the dimensionality reduction model, mapping quality metrics should be low. In our hands, mapping quality metrics failed to distinguish ground-truth OOR cells (Suppl. figure 5A) or neighbourhoods with high fraction of OOR cells (Suppl. figure 5B). When using an ACR design, we could increase the sensitivity by prioritising neighbourhoods where the mapping quality was noticeably worse for the disease dataset cells compared to the control dataset cells (suppl. figure 5C). However, overall we found that using mapping quality metrics to identify OOR populations performed significantly worse than using the logFC estimated from differential analysis.

We tested whether using the healthy atlas in the ACR design reduces the number of control donors needed for detection of disease-specific states. Specifically, we repeated the simulations contrasting cells in a pseudo-disease dataset of 7 donors with a control dataset of increasing numbers of donors (from 3 to 12). We found that across simulations with different OOR cell populations, the true positive rate and especially the false discovery rate varied with the number of control donors with the CR design (Figure 2E). In contrast, OOR-detection outcomes were only marginally affected by the number of control donors when the ACR reference design was used, suggesting that the ACR design can minimise the number of independent control samples required. We also tested how OOR state detection is affected by the size of the atlas dataset used. We expect that if certain normal cell states are undersampled in the atlas dataset, these states might be wrongly predicted to be out-of-reference. We measured the rate of false positives with AR and ACR design when including an increasing number of PBMC studies in the atlas dataset, from 1 to 12 studies, ordered by number of cells in the study (see Methods). While for both designs sensitivity was significantly decreased when using just 1 or 2 studies in the atlas dataset, we observed that the false positive rate increased with smaller atlas datasets with AR design. Conversely almost no false positives were called with ACR design, even with the smallest atlas dataset (suppl. figure 6).

In summary, combining the use of an atlas dataset and of a control dataset for identification of disease-specific cell states significantly reduced the rate of false discoveries compared to using only an atlas or only a control dataset, and led to robust detection of putative disease states, even with a varying number of samples in the control or the atlas dataset.

### Matched controls allow the most sensitive out-of-reference detection

Our results indicate that mapping matched pseudo-disease and control datasets to an atlas dataset enhances the sensitivity to detect OOR states. However in practice there are cases where collecting matched control samples is especially challenging (e.g. brain tumour studies, or studies of osteosarcoma). Therefore, we tested whether selecting a subset of samples from the atlas to use as control dataset in ACR design performed as well as using matched control samples. Specifically, we run the disease-state detection workflow with ACR design defining as control dataset either (a) donors from the same study (matched control), (b) a random subsample of donors from the atlas dataset (random control) or (c) a subset of donors selected by similarity in cell abundances to the disease dataset donors (close control) (suppl. figure 7A, see Methods). In all cases, the control donors are excluded from the atlas dataset used for scVI training. We found that using the matched control always outperformed subsampling from the atlas, and using the close control only marginally reduced the FDR compared to random subsampling (suppl. figure 7B-C). While using any type of control always outperformed differential analysis on the full atlas dataset, these results highlight that comparison against a set of matched controls is always advantageous, likely minimizing false positives driven by hidden confounding factors.

### Mapping to a reference cell atlas improves detection of IFN-stimulated states in COVID-19 patients

We next assessed the benefit of using a healthy atlas for identification of altered states in a real disease cohort. Specifically, we applied our workflow to detect immune cells responding to viral infection in blood from COVID-19 patients. We used a published dataset of single-cell transcriptomes from PBMCs from 90 patients with varying severities of COVID-19 and 23 healthy volunteers (Stephenson *et al*., 2021). As atlas dataset, we used harmonised scRNA-seq profiles from 12 studies that collected PBMCs from a total of 1219 healthy individuals (figure 3A). We applied our workflow to identify disease-associated states, using the healthy PBMC atlas for joint latent embedding (ACR design) or using only the COVID-19 and control datasets (CR design). Also in this case, the atlas dataset is used for scVI model training, but only model weights are used for mapping with scArches and all downstream analysis is performed solely on COVID-19 and control datasets.

To quantify the ability of different reference designs to identify disease-associated states, we tested whether amongst the COVID-19-enriched neighborhoods we could detect cells with activated interferon (IFN) signalling, which is a key pathway involved in antiviral response and a recognized hallmark of COVID-19. We define IFN-stimulated cells by expression of the gene signature defined by (Yoshida *et al*., 2022) (figure 3B, see Methods).

We observed a stronger correlation between the DA logFC and the mean IFN response signature when using the ACR design compared to just using the control as reference (ACR pearson R = 0.63, CR pearson R = 0.51, Fisher’s z transformation p-value < 2.2e−16) (figure 3C). Stratifying by cell type, the correlation is especially strong in the myeloid compartment, where the strongest IFN-stimulation is observed (suppl. figure 8A). Amongst the IFN-low states identified as enriched in COVID-19 with the ACR design, we found primarily plasmablasts and plasma cells (suppl. figure 8B), which are expected to expand in response to COVID-19 (Sette and Crotty, 2021). The CR design additionally detected a strong enrichment in COVID-19 cells in neighbourhoods of IFN-low naive B and T cells, which is in contrast with the widely reported lymphopenia in COVID-19 patients (Chen and John Wherry, 2020), and this enrichment is not explained by expression of proliferation genes (suppl. figure 8C). We also compared reference designs on the ability to distinguish disease-specific phenotypes by clusters and similarity in the latent space: both designs performed equally well in the separation of fine-grained cell types annotated by the original authors (suppl. figure 9A). However, looking into cell annotations where classification performance was different between the two designs, we found that using the ACR design led to more accurate distinction of cell states with known disease-specific phenotypes, including proliferating CD8+T cells, which over-express exhaustion markers, early CD38+ HSCs, exhausted B cells and, notably, a small subset of malignant B cells originating from a COVID-19 patient diagnosed with leukaemia (suppl. figure 9B) (Stephenson *et al*., 2021).

Of note, the number of cells in the control dataset (used for scVI model training with the CR design) is about 3 times smaller than the number of cells in the query dataset. Since Lotfollahi et al. reported robust TL performance with equal sized reference and query datasets, we asked whether using TL might be a suboptimal choice for joint latent embedding with CR design on this dataset. To test this, we compared the disease-state detection workflow using TL with scArches to *de novo* integration, using scVI on the concatenated reference and query datasets. We found that regardless of the latent embedding model used, the integration leveraging the healthy reference atlas leads to more precise detection of IFN-stimulated states (figure 3D). Notably, when using the ACR design, transfer learning performs as well as *de novo* integration. This shows that transfer learning is an effective integration strategy for disease-state identification, and supports the practice of model sharing to overcome the practical hurdles of human data sharing and to reduce the computational burden of atlas-based analysis.

With precise distinction of cell states enriched in COVID-19, we can examine the phenotypic heterogeneity within disease-associated states. In many case-control scRNA-seq studies, cell subtypes are distinguished by iterative rounds of dataset subsetting and subclustering, then differential expression and differential abundance analysis is used to characterise whether subtypes are associated with disease. In blood COVID-19 scRNA-seq studies this procedure frequently leads to a split between an IFN-stimulated COVID-19-associated subcluster and a IFN-low subtype enriched in healthy controls in several PBMC compartments (Ren *et al*., 2021; Yoshida *et al*., 2022). However, IFN-activation is not global, and transitional or alternative pathological phenotypes might be present in PBMCs from COVID-19 patients. In our neighbourhood-level analysis with ACR design, we observe a strong enrichment in cells that express high IFN signature. However, neighbourhoods with relatively low IFN signature are significantly associated with disease, for example amongst classical (CD14+) monocytes (figure 3E), which are notably expanded in the blood of COVID-19 patients. Based on the assignment of cells to neighbourhoods, we separate 3 phenotypes of CD14+ monocytes: normal classical monocytes, COVID-associated IFN-low monocytes and COVID-associated IFN-high monocytes (figure 3F). We observed that the proportion of CD14+ monocytes of different phenotypes changed significantly in patients with different disease severity: the IFN-high state was most prominent in mild and asymptomatic cases, while the IFN-low state became predominant in patients with moderate, severe and critical disease (figure 3G). This observation is in line with the notion that interferon stimulation acts as a protective pathway in the acute phase of infection (Hadjadj *et al*., 2020). In contrast, when defining IFN-high and IFN-low states after differential analysis with CR design, we found the distinction between severity status to be significantly less pronounced (suppl. figure 10A-C). In particular, we find a high fraction of IFN-low COVID-enriched monocytes also in healthy and asymptomatic individuals, indicating that this design wasn’t able to distinguish IFN-low normal monocyte cells from the IFN-low phenotype in severe COVID-19. To characterise the distinct COVID-19 specific phenotypes, we performed differential expression analysis between IFN-high and IFN-low COVID-associated monocytes. Together with IFN-associated genes, IFN-high monocytes showed higher expression of HLA genes, leukocyte-recruiting chemokines (CCL8, CXCL10, CXCL11) and markers of activation (FCGR3A) (suppl. figure 10D, suppl. table 2). In contrast, the IFN-low cells enriched in severe disease over-express S100 genes, previously identified as key markers of COVID-19 severity (Ren *et al*., 2021, Singh and Ali, 2022). This HLA-DRlo S100hi phenotype corresponds to a subset of dysfunctional monocytes associated with severe COVID-19, previously described in an independent cohort through direct comparison of mild and severe cases (Schulte-Schrepping *et al*., 2020).

In summary, using atlas and control reference on a COVID-19 cohort led to precise detection of blood cell states associated with response to infection, without requiring subclustering analysis, while still distinguishing phenotypically distinct pathological cell states.

## Discussion

In this study we quantitatively examine the impact of the choice of reference dataset on the task of identification of altered cell states from scRNA-seq data of diseased tissues. Currently, single-cell genomics datasets on patient cohorts include data from tens of individuals, processed with uniform experimental protocols. Many disease studies collect matched controls alongside the disease samples, with similar demographic and experimental protocol characteristics (e.g. Nehar-Belaid *et al*., 2020; Schulte-Schrepping *et al*., 2020; Stephenson *et al*., 2021; R. K. Perez *et al*., 2022; Yoshida *et al*., 2022). In parallel, the single-cell community is moving towards the generation of large, harmonised collections of data from the same tissue, with the explicit aim to serve as maps for contextualization of altered cellular conditions, and several studies use these datasets as references for discovery of disease-states, also in the absence of controls (Guo *et al*., 2020; Jardine *et al*., 2021; Szabo *et al*., 2021; Reichart *et al*., 2022; Sikkema *et al*., 2022). The option to use large curated datasets is amenable to efficient data re-use and especially for tissues where collection of matched healthy controls is challenging, such as studies of the brain (Olah *et al*., 2020; Leng *et al*., 2021). In this work we demonstrate on simulations and real disease datasets that atlas datasets are not a substitute for control samples, but they should be used to increase the sensitivity and precision of disease-state discovery.

We present a quantitative benchmark for the task of detection of out-of-reference cell states. We used these *in silico* experiments to compare reference selection choices, but also to highlight shortcomings of mapping quality-control metrics that have been used as heuristics to prioritise disease-specific cell states (suppl. figure 5). We designed evaluation experiments and chose methods for integration and differential analysis with the specific use-case of disease datasets in mind. We believe our results will extrapolate beyond to other types of case-control studies, as long as certain assumptions apply: we assume that all the cell states observed in the control dataset are also found in the atlas dataset and that only a fraction of cell types are altered in the disease datasets. We believe these are reasonable assumptions when comparing healthy and pathological states in the same human tissue. However, cellular phenotypes might be different with alternative case-control designs, for example when measuring the effect of genetic perturbations in cell lines, where we expect less diversity of cellular phenotypes in the wild-type control and all perturbed cells might be affected to an extent (Replogle *et al*., 2022).

With the disease analysis scenario in mind, we use a transfer learning (TL) method for joint dimensionality reduction (Lotfollahi *et al*., 2022), as an efficient alternative to integration of the concatenated disease and reference datasets. We made this choice envisioning that in the future the single-cell community would be sharing trained models for fast re-use or integrated atlases, which do not require downloading and reprocessing data from millions of cells. We expect our reference design analysis results to be robust to the use of *de novo* integration, as shown on the COVID-19 application (figure 3D), or to other TL-based methods for query-to-reference mapping (Hao *et al*., 2021; Kang *et al*., 2021). To date, these and other integration methods have been primarily benchmarked on the tasks of batch correction, automatic cell type annotation and fast analysis of new data generated from the same biological condition (Tran *et al*., 2020; Chazarra-Gil *et al*., 2021; Luecken *et al*., 2022). While the systematic comparison of different integration methods on disease datasets is beyond the scope of this study, the benchmarking set-up and python package presented in this study (https://github.com/MarioniLab/oor_benchmark) could serve as groundwork for future comparisons focused on alternative integration methods.

The results from our analyses have important implications for experimental design of single-cell genomics studies of diseased tissues. We show that, when an atlas dataset is available, it is possible to reduce the number of control samples without introducing false discoveries and minimal impact on sensitivity (figure 2E). In addition, our analyses indicate that matched controls are always to be preferred over selecting a subset of samples from published data to use as control (suppl. figure 7). To an extent, accounting for confounding covariates in differential analysis based on available metadata might reduce the false discovery rate when no matched controls are available. However, it is to be expected that comparing uniformly processed healthy and disease samples from a similar population will minimise variability associated with known and hidden confounders. Selecting samples from published data based on phenotypic similarity led to a small improvement in performance compared to selection of samples at random. While here we applied a naive approach to define sample similarity (suppl. figure 7A), novel methods to model sample-level heterogeneity in scRNA-seq data are being explored (Chen *et al*., 2020; Boyeau *et al*., 2022; Mitchel *et al*., 2022). These could improve the matching of disease samples to optimal controls, and provide new insights into which technical and demographic variables are likely to affect disease-to-healthy comparisons.

Having demonstrated the advantages of atlas-based analysis of disease and control cohorts, this work emphasises the importance of meta-analysis to build population-scale cell atlases of human tissues. For certain tissues the wealth of available data has reached the scale of cell atlases, such as for blood, lung (Sikkema *et al*., 2022), heart (Litviňuková *et al*., 2020; Hocker *et al*., 2021; Koenig *et al*., 2022) or gastro-intestinal tract (Elmentaite *et al*., 2021). However, efforts for integration and harmonisation are still in progress and we expect these integrated datasets to be frequently updated, to incorporate data from more individuals. While this process is underway, false positives might arise in atlas-based analysis of disease datasets if normal cell states are missed in the atlas dataset because of insufficient sampling. We show that joint mapping of disease and control samples can help to maintain a low rate of false positives with smaller atlas datasets (suppl. figure 6), making disease analysis more robust to atlas updates. We hope this encourages early sharing of beta versions of integrated tissue atlases (as exemplified by the Azimuth developers https://azimuth.hubmapconsortium.org/), as these could already provide a valuable resource for the study of human pathologies.

Our analysis on a COVID-19 cohort highlights how using a cell atlas to contextualise data from a case-control study leads to more sensitive detection of cell states associated with infection (figure 3). While the simulated OOR cases exemplify a case where a single cluster of transcriptionally distinct cells are disease-specific, the activation of the interferon signalling in response to infection represents an example of a common response across cell types, which is rarely captured as a principal source of variation, when performing a cluster-centric analysis. With precise distinction of cell states enriched in COVID19, we can examine the phenotypic heterogeneity within disease-associated states. In many case-control scRNA-seq studies, cells with the same annotation from the disease and healthy condition are contrasted with DE analysis, but this procedure might dilute the disease-specific signature if diseased tissue contains cells with both the normal phenotype and the pathological phenotype, or several distinct pathological subtypes. Alternatively, distinct cell subtypes are distinguished by iterative rounds of dataset subsetting and subclustering, then differential abundance analysis is used to characterise whether subtypes are associated with disease. In blood COVID-19 scRNA-seq studies, this procedure leads to a binary distinction between an IFN-high COVID-19-associated subcluster and an IFN-low subtype enriched in healthy controls (Yoshida *et al*., 2022). With subpopulation-level differential abundance analysis (Dann *et al*., 2022) we capture heterogeneous COVID-19 enriched states in several cell types, including CD14+ monocytes (figure 3E-G). While the original study reported relative increase in IFN-activated CD14+ monocyte abundance with COVID-19, we identify distinct IFN-high and IFN-low pathological subtypes, associated with distinct disease severities. These subtypes correspond to subsets of dysfunctional monocytes previously described in an independent cohort, through explicit comparison of mild and severe cases (suppl. figure 10D) (Schulte-Schrepping *et al*., 2020). This example demonstrates that precise disease-state detection with ACR design enables the study of transitional and heterogeneous pathological cell states.

In conclusion, we demonstrate that the combined use of a cell atlas and matched controls as references enables the most precise identification of affected cell states in disease scRNA-seq datasets. We envision our analysis will instruct the design of new cohort studies, guide efficient data re-use and provide guiding principles for analysis of disease dataset and construction of cell atlases.

## Supporting information

Metadata on PBMC studies

DE analysis results on pathological CD14+ monocytes

## Acknowledgements

We thank M. Morgan and R. Lindeboom for critical reading of the manuscript; K. Meyer, R. Elmentaite, A. Missarova and all members of the Marioni lab and Teichmann lab for valuable discussions and feedback on this project. The PBMC studies included in this work were selected using the materials from the Chan-Zuckerberg Initiative workshop on “Assembling Tissue References ”, which were kindly shared by L. Dratva. Figure 1A was created with BioRender.com.

## Contributions

E.D. and J.C.M. conceptualized this study. E.D. wrote the benchmarking package and performed the analysis. All authors interpreted the results, wrote and approved the manuscript. J.C.M. oversaw the project.

## Methods

### PBMC data preprocessing

We collected raw gene expression counts and cell type annotations from healthy peripheral blood mononuclear cell (PBMC) 10X Genomics scRNA-seq data from 13 publications, available via the cellxgene portal (https://cellxgene.cziscience.com/collections) (Suppl. Table 1). During harmonisation, we sampled 500 cells for each sample to reduce the computational burden of this analysis, while maintaining sample-level diversity, and we excluded samples for which less than 500 cells were detected, retaining in total 1268 samples from 1248 individuals. We subsequently filtered cells where at least 1000 mRNA molecules were detected and genes that were expressed in at least one cell. This resulted in a dataset of 599379 high-quality cells.

To generate a unified cell type annotation, we integrated all normal cells from different studies in a common latent space using the scVI model, as implemented in the python packages scvi-tools (Lopez *et al*., 2018; Gayoso *et al*., 2022). Briefly, we selected the 5000 most highly variable genes (HVGs) based on dispersion of log-normalised counts, as implemented in scanpy. We trained the scVI model on raw counts, subsetting to HVGs, considering the library ID as batch (model parameters: `n_latent = 30, gene_likelihood = ‘nb’, use_layer_norm = “both”, use_batch_norm =` “n`one”, encode_covariates = True, dropout_rate = 0.2, n_layers = 2;` training parameters: `early_stopping=True,train_size=0.9, early_stopping_patience=45, max_epochs=200, batch_size=1024,limit_train_batches=20`). We constructed a k-nearest neighbor graph based on similarity in the scVI latent dimensions, using k=50. Cells were clustered using the leiden algorithm with resolution=1.5. Subsequently, clusters were annotated by majority voting using the harmonised cell type labels available via cellxgene. During this process, one cluster of cells was excluded as potentially containing doublets. After this final filtering the dataset included 597321 cells annotated into 16 cell types.

### Simulation experiments

#### Data splitting into atlas/control/perturbation

To simulate the attributes of disease, atlas and control datasets, we select donors from one study (query study, 29 healthy donors, Stephenson et al. 2021) and we split these at random with equal probabilities into a disease subset (16 donors) and a control subset (13 donors). The data from the remaining 12 studies comprises the atlas dataset (1219 donors). To simulate the presence of an out-of-reference cell state, we select one cell type label and remove all cells with that label from the control and atlas dataset. We repeat this simulation with 15 annotated cell types in the PBMC dataset (neutrophils were excluded, as they were under-represented in the Stephenson study).

#### Latent space embedding

For each simulated atlas/control/disease dataset assignment, we embed the reference and query datasets into a common latent space using transfer learning with scArches (Lotfollahi *et al*., 2022) on scVI models (Lopez *et al*., 2018), using the implementation in the python package scvi-tools v0.17.4 (Gayoso *et al*., 2022). Briefly, we selected the 5000 most highly variable genes (HVGs) in the reference dataset based on dispersion of log-normalised counts, as implemented in scanpy. We trained the scVI model on raw counts of the reference dataset, subsetting to HVGs, considering the sample ID as batch and specifying the recommended parameters to enable scArches mapping (use_layer_norm = “both”, use_batch_norm = “none”, encode_covariates = True, dropout_rate = 0.2, n_layers = 2). Models were trained for 400 epochs or until convergence. Next, we perform transfer learning on the query dataset(s) from the model trained on the reference, running the model for 200 epochs and setting the weight_decay parameter to 0. Reference and query dataset for latent space embedding are defined as follows for the three reference designs:

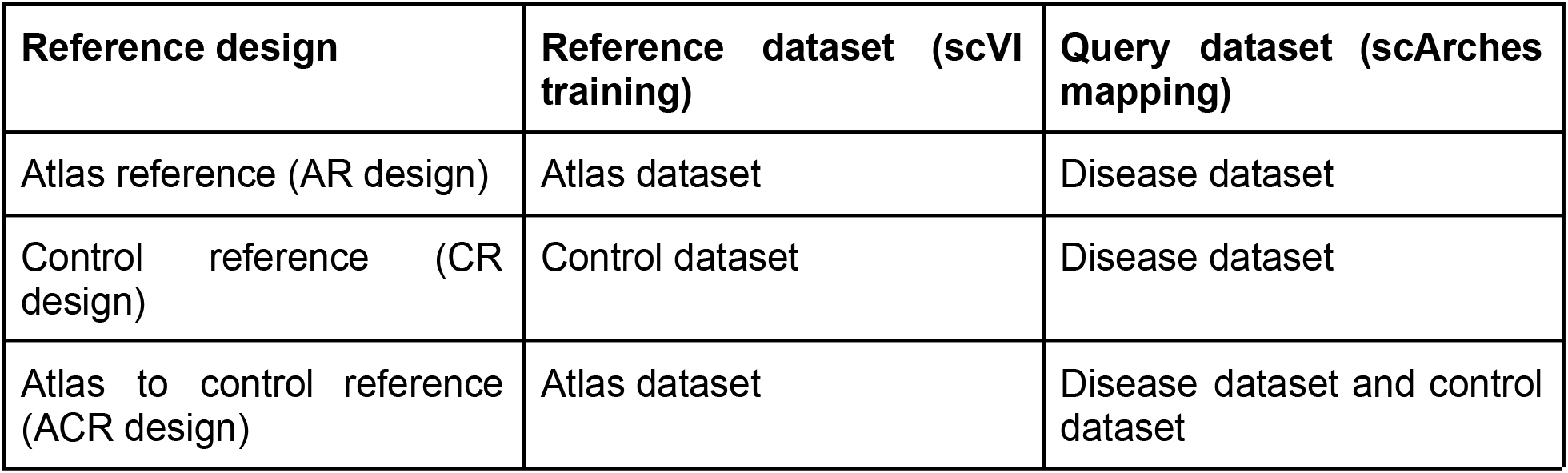

#### Differential abundance analysis with Milo

To find cell states enriched in the disease dataset we use the Milo framework for differential abundance analysis on cell neighbourhoods (Dann *et al*., 2022) using the implementation in the package milopy (https://github.com/emdann/milopy, v0.1.0). Briefly, we compute the k-nearest neighbor (KNN) graph of cells in the reference and disease dataset based on the latent embedding. The reference datasets for differential analysis are defined as follows for the three reference designs:

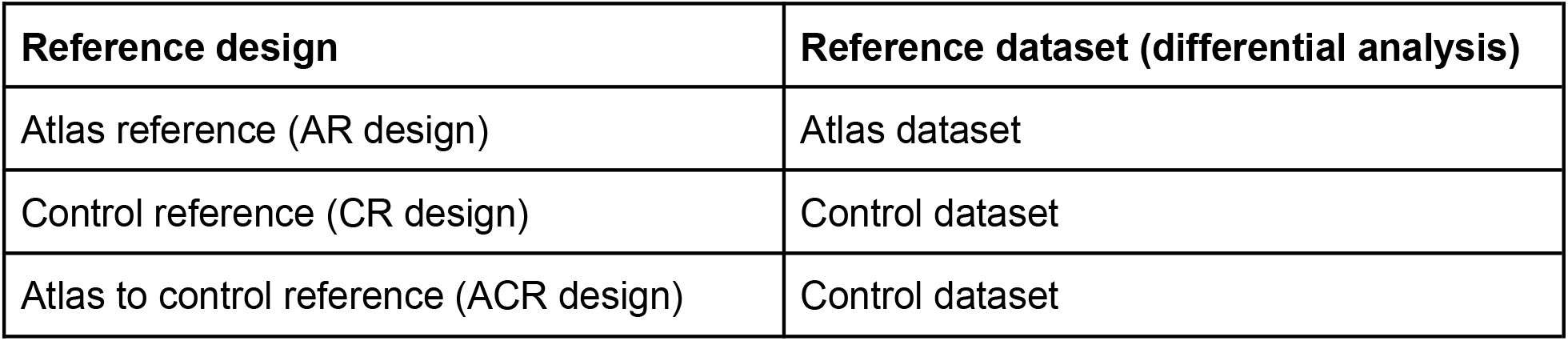

| Reference design | Reference dataset (differential analysis) |
| --- | --- |
| Atlas reference (AR design) | Atlas dataset |
| Control reference (CR design) | Control dataset |
| Atlas to control reference (ACR design) | Control dataset |

Of note, for the ACR design the atlas dataset is not considered when constructing the KNN graph. We set the value of k to be the equal to the total number of samples times 5, up to a maximum of k = 200 (this upper limit is set for memory efficiency reasons), following the indications in Dann et al. We assigned cells to neighborhoods (milopy.core.make_nhoods, parameters: prop = 0.1) and counted the number of cells belonging to each sample in each neighborhood (milopy.core.count_cells). We assigned to each neighborhood a cell type label based on majority voting of the cells belonging to that neighborhood. To test for enrichment of cells from the disease dataset, we model the cell count in neighborhoods as a negative binomial generalised linear model, using a log-linear model to model the effects of disease status on cell counts (log-Fold Change, logFC). Although the split between control and disease samples was balanced in terms of available metadata, in the query study there is a known batch effect between the three sites from which samples were collected (Stephenson *et al*., 2021). Therefore we included the site identity as a confounding covariate in the differential abundance model when using the ACR and CR design, although we found the results presented here were robust even without modelling this confounder (data not shown). We control for multiple testing using the weighted Benjamini-Hochberg correction as described in Dann et al. (SpatialFDR correction). Unless otherwise specified, neighbourhoods were considered enriched in disease cells if the SpatialFDR < 0.1 and logFC > 0.

#### Sensitivity analysis

For each simulation (i.e. with different OOR cell state and reference design) we define a neighbourhood as an OOR state (true positive) if the percentage of OOR cells in the neighbourhood is more than 20% of the maximum percentage observed in that simulation. This threshold selection aims to quantify the ability to detect the neighbourhoods where the largest number of OOR cells is found, even when the atlas dataset is included in the KNN graph (AR design), and the majority of cells in neighbourhoods always belong to the atlas dataset. The selected thresholds for each experiment are shown in suppl. figure 1. We calculate true positive rates, false positive rates and false discovery rates considering neighborhoods where the SpatialFDR < 0.1 and logFC > 0 as predicted positives.

With precision-recall curve analysis, we quantify the ability to detect true positive OOR states with different thresholds of logFC, without considering the significance estimated with the SpatialFDR, using the implementation in scikit-learn (Pedregosa, Varoquaux and Gramfort, 2011).

#### Mapping quality control metrics

We compare quality control (QC) metrics for scArches on the mapping of disease dataset onto the atlas dataset (mapping QC score), quantifying whether OOR cells and cell states are detectable by low QC score (i.e. higher uncertainty during query-to-reference mapping). For the label transfer uncertainty, we use the method presented by (Lotfollahi *et al*., 2022). Briefly, we train a weighted KNN classifier for cell type labels on the latent embedding of the scVI reference dataset, setting the number of neighbors to k=100. For each cell in the query dataset, we extract its k nearest neighbors in the reference and the corresponding Euclidean distances, adjusted by a Gaussian kernel. We compute the probability *P*(*y*|*c*) of assigning each label *y* to the query cell *c* by normalising across all adjusted distances. The label uncertainty corresponds to 1 − *max*_y_ (*P*(*y*|*c*)).

For the reconstruction error metric, we take *s* = 50 samples from the posterior of the scArches model to generate profiles of predicted gene expression counts 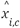for the highly variable genes used in training. We define the reconstructed gene expression profile 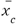 for each query cell *c* as:

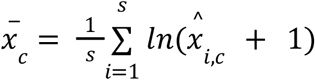

Given the true log-normalised gene expression profile for each cell *x*_*c*_, we define the reconstruction error as the cosine distance between *x*_*c*_ and 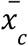.

For both label uncertainty and reconstruction error, higher values correspond to lower mapping quality. To compare the mapping QC metrics to the differential abundance logFC on neighbourhoods defined with the ACR design (suppl. figure 5B) we calculate the average mapping QC score over all disease cells in the same neighbourhood. For the comparison between mapping QC scores in control and disease cells with ACR design (suppl. figure 5C), for each neighbourhood we compute the difference between the average QC scores over disease cells and over control cells.

#### Control and atlas size analysis

For the analysis with varying number of control donors (figure 2E), we selected the simulations with the 5 OOR cell populations that had the highest average TPR between CR and ACR design in the previous analysis (figure 2D). For each simulation, we selected the 7 donors from the disease dataset that had the highest fraction of cells in the OOR cell population. Subsequently, we selected a random subset of *n* donors (with 3 < *n* < 12) from the control dataset and performed disease-state identification with the CR or ACR design, as described above. For each *n*, we repeated the simulation with 5 different initializations of the control donor selection.

#### Atlas subsampling analysis

To assess whether a shallow atlas dataset introduces false discoveries in disease-state identification (suppl. figure 6), we used all 29 donors from the query dataset in disease and control datasets, and subsampled the atlas dataset removing data from 1 to 11 studies (ordering studies by total number of cells), and performed disease-state identification with AR and ACR design.

#### Control selection from atlas sampling analysis

We tested whether selecting a subset of samples from the atlas dataset to use as control samples for disease-state identification with ACR design performs comparably to using matched controls. We applied the following unsupervised selection strategy, based on similarity between cell state proportions (suppl. figure 7A): we take latent dimensions learnt by training scVI on the atlas dataset and mapping the disease dataset with scArches (i.e. as described above for AR design). In this latent space, we construct Milo neighbourhoods and the matrix of cell counts of dimension samples x neighbourhoods, as described in the section “Differential abundance analysis with Milo”. We log-normalise cell counts per sample (running scanpy.pp.normalize_total with parameters target_sum=10000, and scanpy.pp.log1p) and perform principal component analysis on the matrix of log-normalised cell counts. This generates a d-dimensional space (d=10) on which we compute Euclidean distances between disease and atlas samples. In this space, for each disease sample we take the closest atlas sample and we consider this set to be the “close controls”. If one atlas sample is the nearest neighbor to multiple disease dataset samples, we look at the second nearest neighbors for these samples and take those with the smallest Euclidean distance.

We rerun the disease state identification workflow with ACR design using as control datasets: (i) matched controls (samples from the same study), (ii) close controls and (iii) randomly selected samples from the atlas dataset. The same number of control samples was used in all three cases.

### Design comparison on COVID dataset

#### Data preprocessing and model training

We downloaded data from COVID-19 and healthy PBMCs from (Stephenson *et al*., 2021), via the cellxgene portal (collection ID: ddfad306-714d-4cc0-9985-d9072820c530). We sampled 500 cells for each sample to reduce the computational burden of this analysis, while maintaining sample-level diversity, and we excluded samples for which less than 500 cells were detected. We excluded cells where less than 1000 mRNA molecules were detected and we excluded data from 3 samples that were profiled with the Smart-seq-2 protocol. As cell type annotations, we use the high-level annotation from the original authors.

As the atlas dataset we used the healthy PBMC data described above, excluding the healthy PBMC profiles from Stephenson et al. 2021. Reference model training and query mapping was performed as described above. After query-mapping, control and COVID-19 cells were embedded in a KNN graph (k=100), which was used to build neighbourhoods and perform differential abundance analysis with Milo as described above. For the comparison of *de novo* integration and query-mapping (figure 3D), scVI training was performed as described above on concatenated atlas, control and COVID-19 datasets (ACR design) or control and COVID-19 datasets (CR design).

#### IFN signature calculation

To define IFN-stimulated cells, we aggregate expression of a set of IFN-associated genes defined by Yoshida et al. 2021 (including BST2, CMPK2, EIF2AK2, EPSTI1, HERC5, IFI35, IFI44L, IFI6, IFIT3, ISG15, LY6E, MX1, MX2, OAS1, OAS2, PARP9, PLSCR1, SAMD9, SAMD9L, SP110, STAT1, TRIM22, UBE2L6, XAF1 and IRF7), using the scanpy function scanpy.tl.score_genes() to quantify signature expression for each cell. A threshold of IFN-signature > 0.05 was used for precision-recall analysis.

#### Label prediction with KNN classifier

To assess the difference between CR and ACR design on the latent space learnt with transfer learning, we trained a k-nearest neighbor classifier to learn the original fine cell type label from Stephenson et al. 2021. We used the implementation in scikit-learn (function: KNeighborsClassifier, parameters: n_neighbors=30, metric=‘euclidean’). The dataset was split randomly into training (80% of cells) and testing sets (20% of cells) and the reported performance was calculated on the test set for 10 different test-train splits (suppl. figure 9).

#### CD14+ monocyte disease-state analysis

For the analysis on COVID-19-associated monocyte subsets, we focused on the neighbourhoods annotated as CD14+ monocytes based on majority voting, as described above. We split CD14+ monocyte neighborhoods into IFN-high COVID-19 neighbourhoods (SpatialFDR < 0.1, logFC > 0 and IFN signature > 0.2), IFN-low COVID-19 neighbourhoods (SpatialFDR < 0.1, logFC > 0 and IFN signature < 0.2) and healthy neighborhoods (remaining neighborhoods). To assign cells to one of these 3 phenotypes, we computed, for each cell, the number of neighborhoods of each phenotype to which that cell belongs (since Milo neighborhoods can be partially overlapping) and we labelled the cell based on the most representative phenotype (if the cell was found in at least 3 neighbourhoods of that phenotype, otherwise the cell was annotated as mixed CD14+ monocyte phenotype).

For differential expression analysis, we aggregated gene expression profiles by summing counts by sample and CD14+ monocyte phenotype and performed differential expression testing using the edgeR quasi-likelihood test (Robinson and Oshlack, 2010) using the implementation in the R package glmGamPoi (Ahlmann-Eltze and Huber, 2021), using 1% FDR (suppl. table 2).

## Code and data availability

The functions for benchmarking out-of-reference state detection, including code for preprocessing, data splitting, integration, differential analysis and evaluation metrics, have been made available as a python package at https://github.com/MarioniLab/oor_benchmark. Notebooks and scripts to reproduce all analysis presented in the manuscript are available at https://github.com/MarioniLab/oor_design_reproducibility.

All the data used for analysis is publicly available (see suppl. table 1). Processed data objects and trained scVI models are available via figshare (https://doi.org/10.6084/m9.figshare.21456645.v1).

## Supplementary Figures

**Supplementary figure 1:**
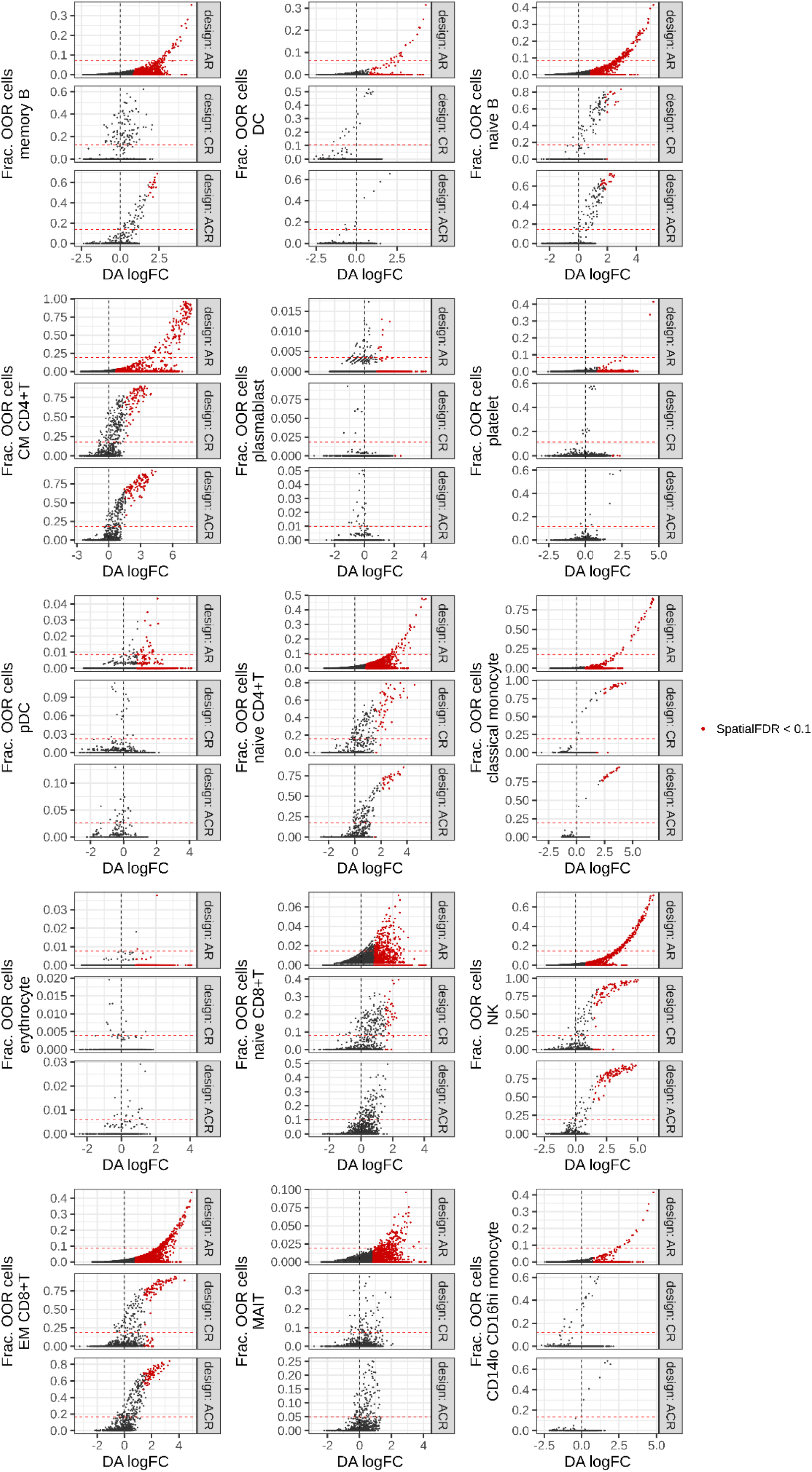
Out-of-reference recovery across simulations. Scatterplot of differential abundance log-Fold Change (DA logFC) against fraction of out-of-reference (OOR) cells for each neighbourhood, in simulations with different unseen cell populations (indicated in y-axis). Colored points indicate neighbourhoods where the enrichment was significant (10% SpatialFDR, logFC > 0). The dotted red line indicates the threshold used to define true positives (20% of the higher fraction in the simulation).

**Supplementary figure 2:**
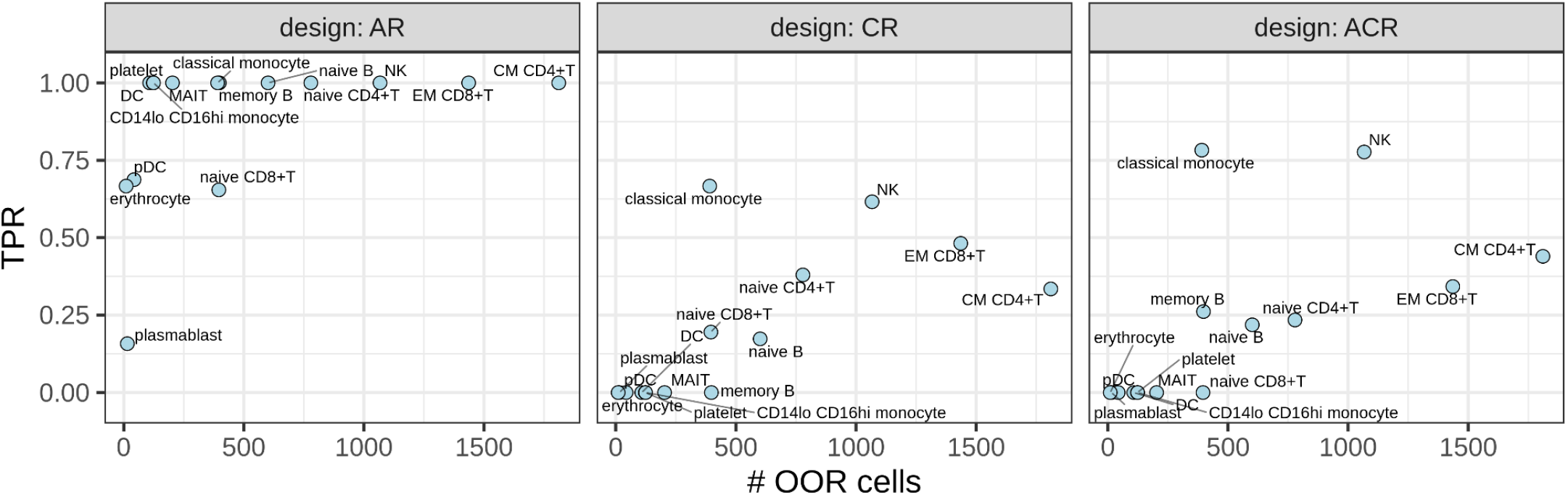
Statistical power is dependent on the size of the OOR cell state across reference designs. Scatterplot of number of cells in the simulated OOR state (x-axis) against the true positive rate (TPR, y-axis) of OOR state detection with alternative reference designs.

**Supplementary figure 3:**
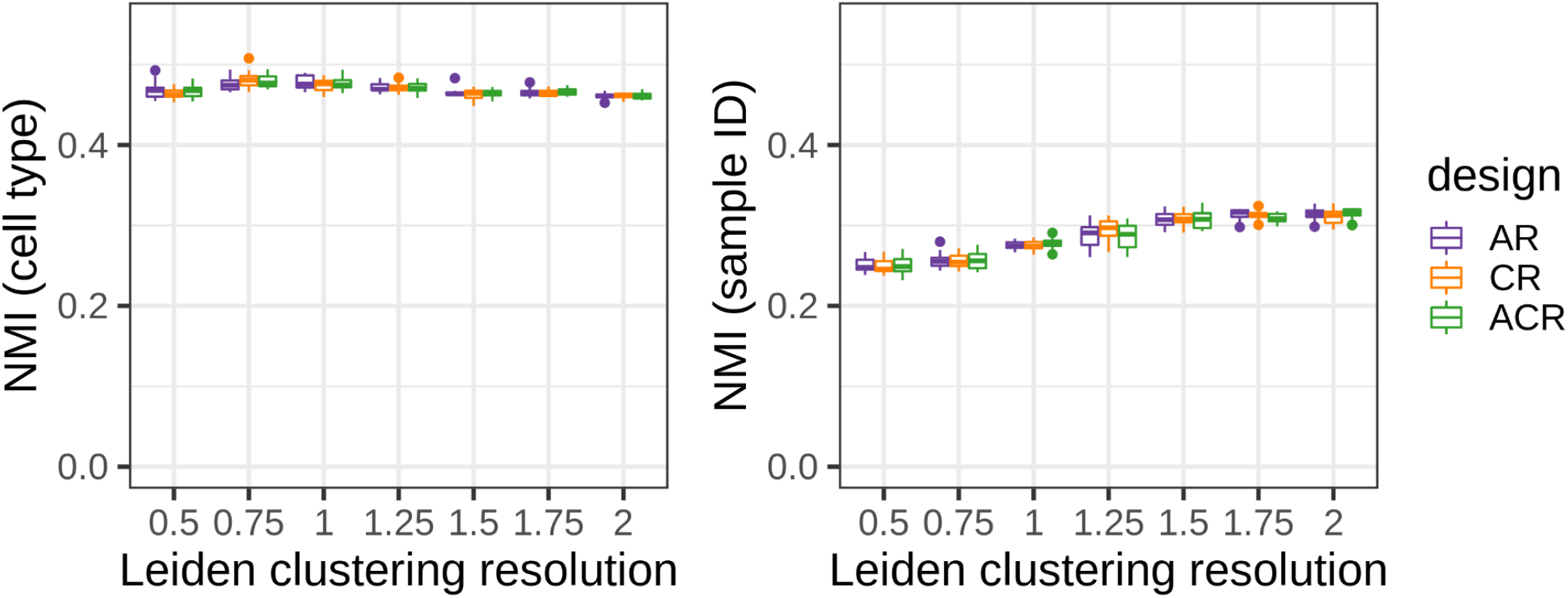
Batch correction and biological conservation is comparable between latent dimensions learnt with different reference design: quantification of overlap between cell type labels (as a measure of biological conservation, left) and sample IDs (as a measure of batch effect, right) and clusters of disease cells on latent dimensions after scArches mapping with different designs (color). The overlap between clusters and covariates is measured by the Normalised Mutual Information (NMI), using the implementation in scikit-learn v1.1.2 (Pedregosa, Varoquaux and Gramfort, 2011). Each box plot shows the median and interquartile range for simulations with different OOR cell populations. NMI values for leiden clustering with increasing resolution (x-axis) are shown.

**Supplementary figure 4:**
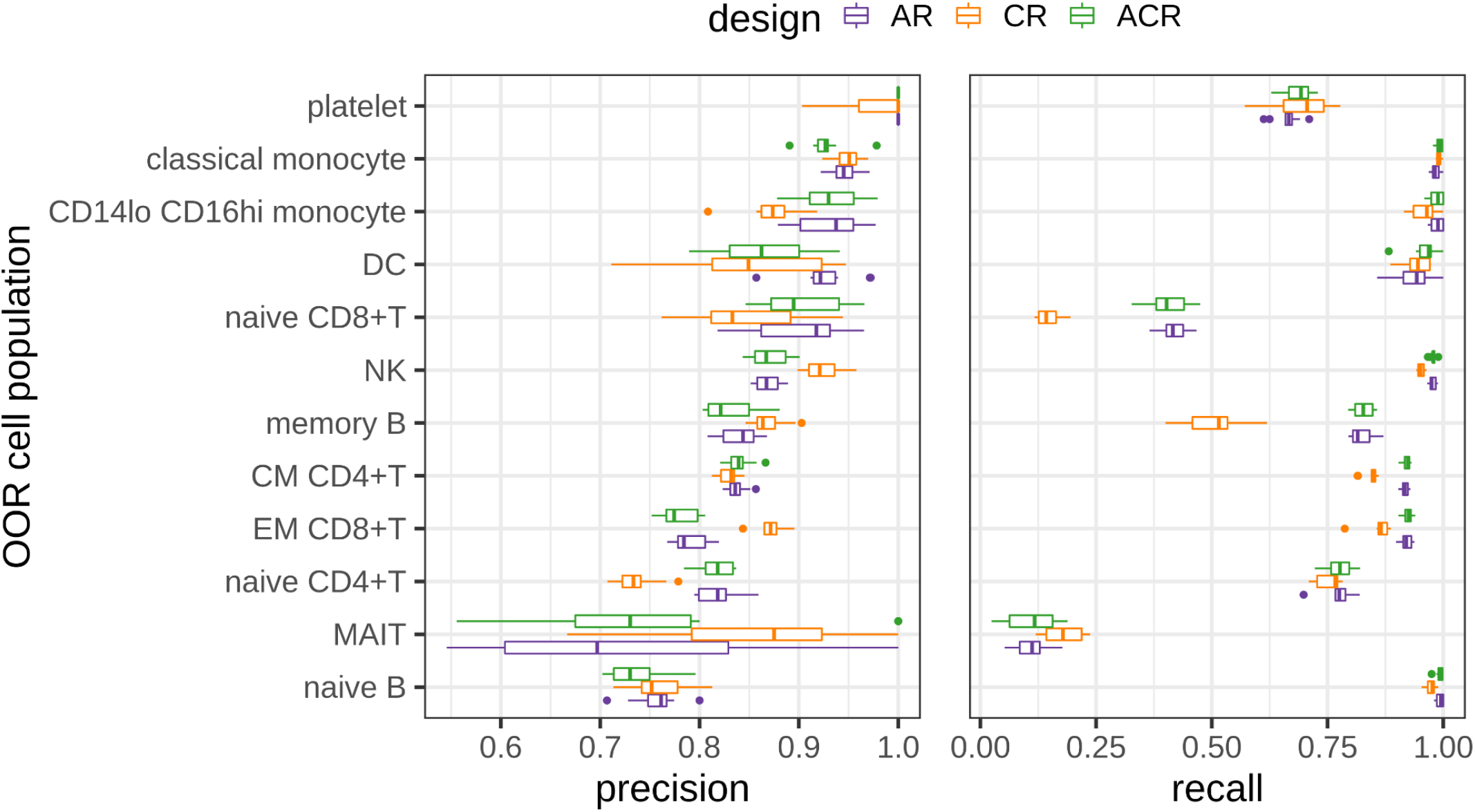
The out-of-reference state is not always distinguished by position in the latent space after scArches mapping. estimated precision (left) and recall (right) for simulations with different OOR cell populations (y-axis) when training a KNN classifier on the latent space of disease cells to distinguish the true OOR cells. Each box plot shows the median and interquartile range for 10 classifiers trained with a different split of cells into training and test set. Precision and recall on the test set are shown. This analysis was performed only on simulations where at least 100 cells were found in the OOR state in the disease dataset.

**Supplementary figure 5:**
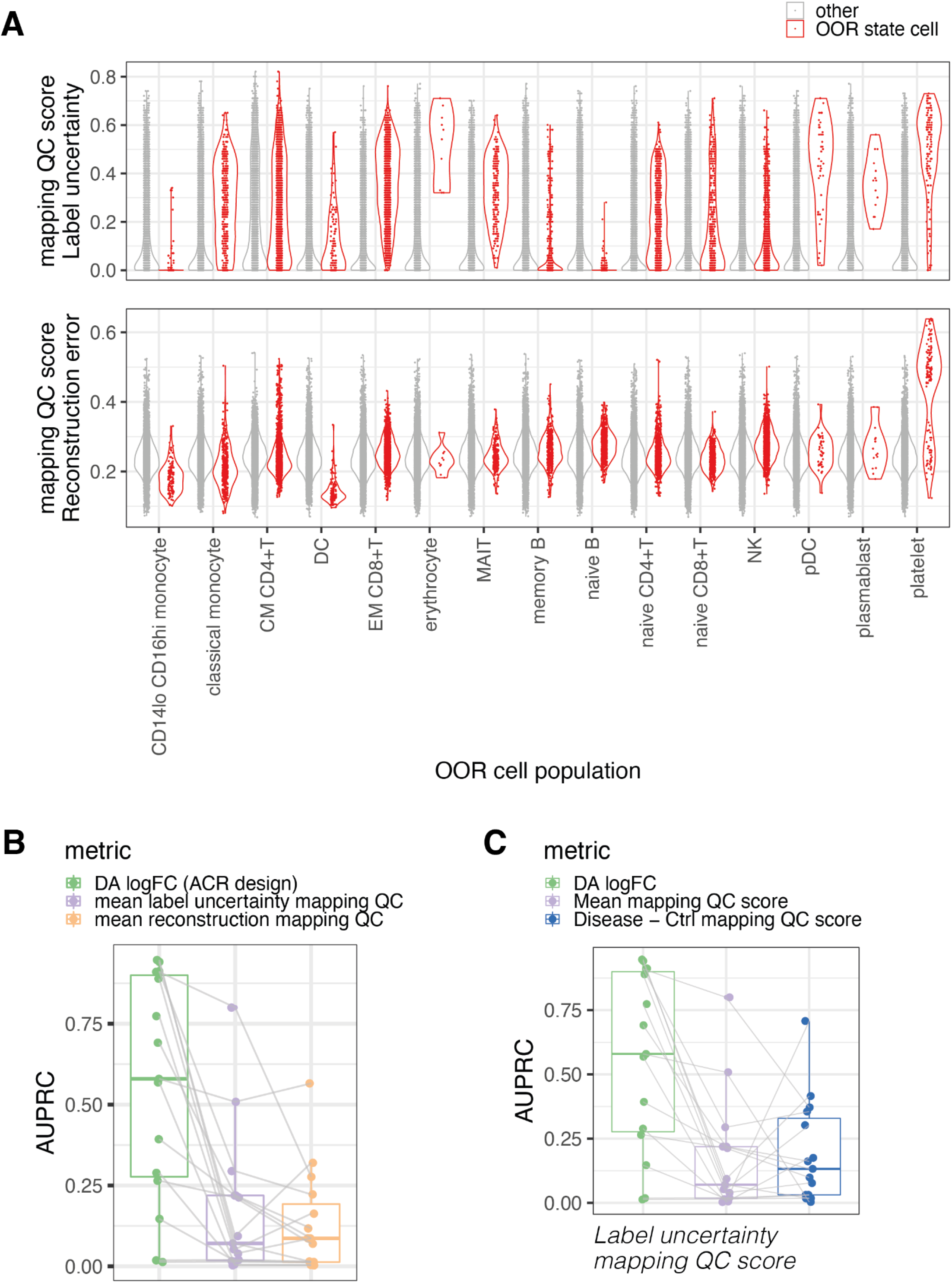
Comparison to mapping quality control metrics. **(A)** Violin plots of mapping QC scores for cells in OOR state (red) and all other cells (grey) across 15 simulations with different OOR cell populations (x-axis). The top plot shows label uncertainty as mapping QC score, the bottom plot shows reconstruction error as mapping QC score. **(B)** Boxplot of Area Under the Precision-Recall Curve (AUPRC) for detection of OOR state neighbourhoods with 3 different metrics: differential abundance log-Fold Change with ACR design (DA logFC, green), mean label uncertainty (purple), mean reconstruction error (orange). Points represent simulations with different OOR populations. Tests on the same simulated data are connected. **(C)** As in (B), but we compare DA logFC and mean label uncertainty to the difference between mean label uncertainty of disease cells and mean label uncertainty of control cells in the same neighbourhood (blue).

**Supplementary figure 6:**
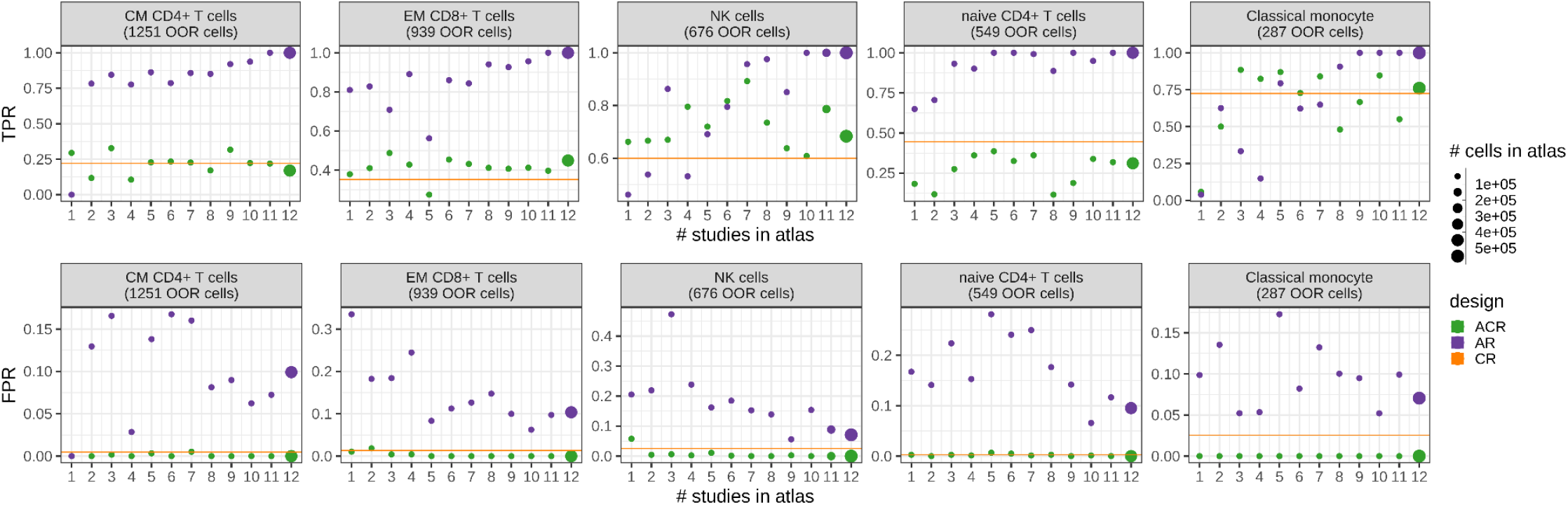
True positives and false positives with decreasing size of atlas dataset. true positive rate (TPR, top y-axis) and false positive rate (FPR, bottom y-axis) of OOR state detection for simulations with an increasing number of studies included in the atlas reference (x-axis) for AR and ACR design (colour). Studies were ordered by total number of cells. Point size is proportional to the number of cells in the atlas dataset. Results from simulations with five different OOR cell states are shown, selected by top mean TPR across designs in (Figure 2D). The orange line denotes the TPR and FPR values for the CR design (i.e. not using an atlas dataset).

**Supplementary figure 7:**
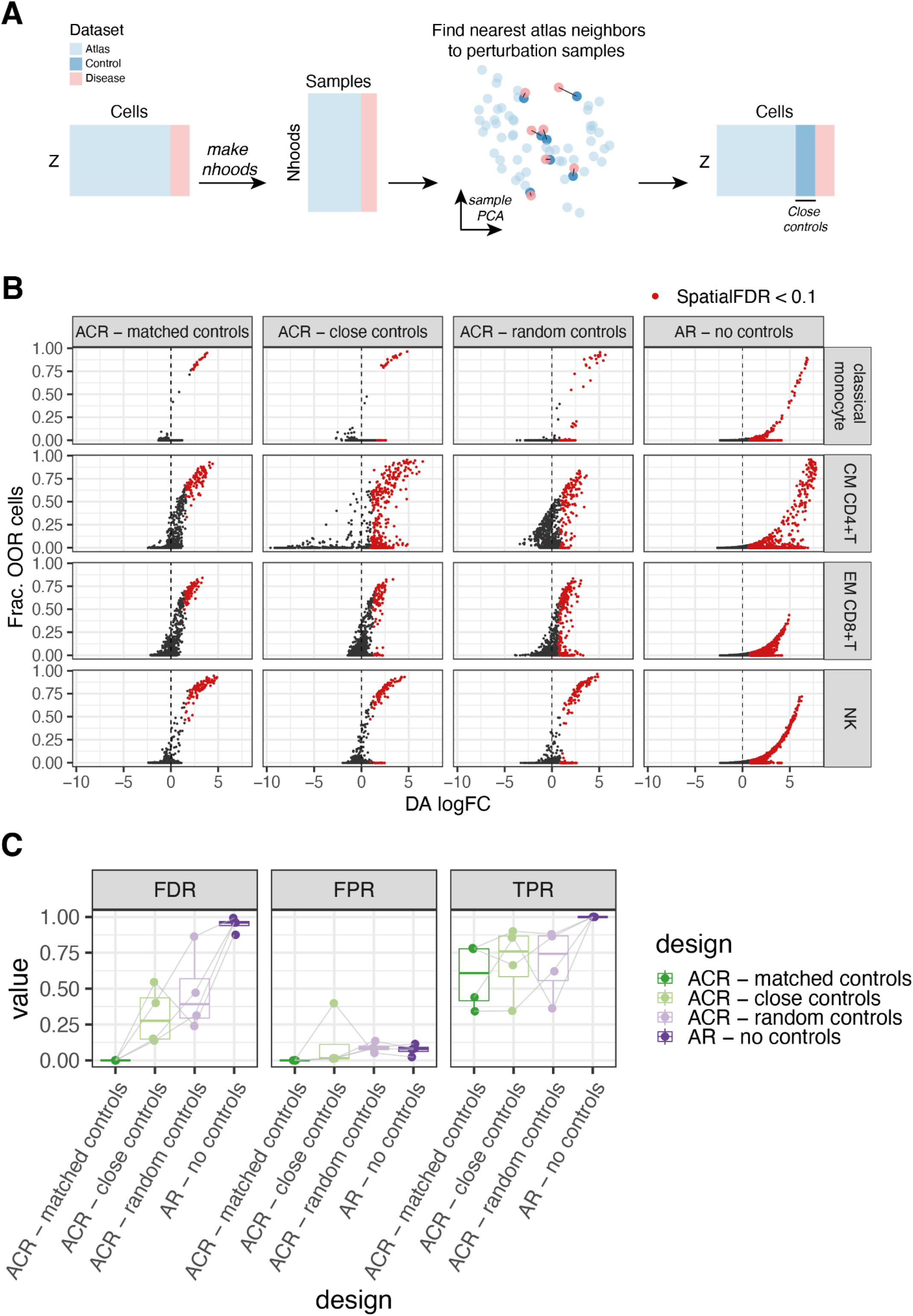
Comparison of matched controls and controls selected from the atlas dataset. **(A)** Schematic of method used for unsupervised selection of “close controls’’ from the atlas samples: the latent space (Z) learnt from mapping of the disease dataset to the atlas dataset was used to construct a KNN graph and neighbourhoods. We generated a count matrix storing the number of cells from each sample in each neighbourhood (as used in Milo differential abundance analysis). We projected samples over principal components (PCs) that explain the highest variance in the neighbourhood count matrix and we computed euclidean distances between disease and atlas samples in the top 10 PCs. For each disease sample we take the closest atlas sample and we consider this set of samples to be the “close control” dataset. **(B)** Scatterplot of differential abundance log-Fold Change (DA logFC) against fraction of out-of-reference (OOR) cells for each neighbourhood, in simulations with different unseen cell populations (rows), with different control samples used for ACR design or no controls (AR design). Colored points indicate neighbourhoods where the enrichment was significant (10% SpatialFDR and logFC > 0). **(C)** Quantitative comparison of performance with different control samples used for ACR design or no controls (AR design) in detection of OOR cell states (neighbourhoods where the fraction of cells from the perturbation-specific state is higher than 20% of the maximum fraction). To compare performance considering the logFC and confidence (10% SpatialFDR), we measured the false discovery rate (FDR), false positive rate (FPR) and true positive rate (TPR). Tests on the same simulated data are connected.

**Supplementary figure 8:**
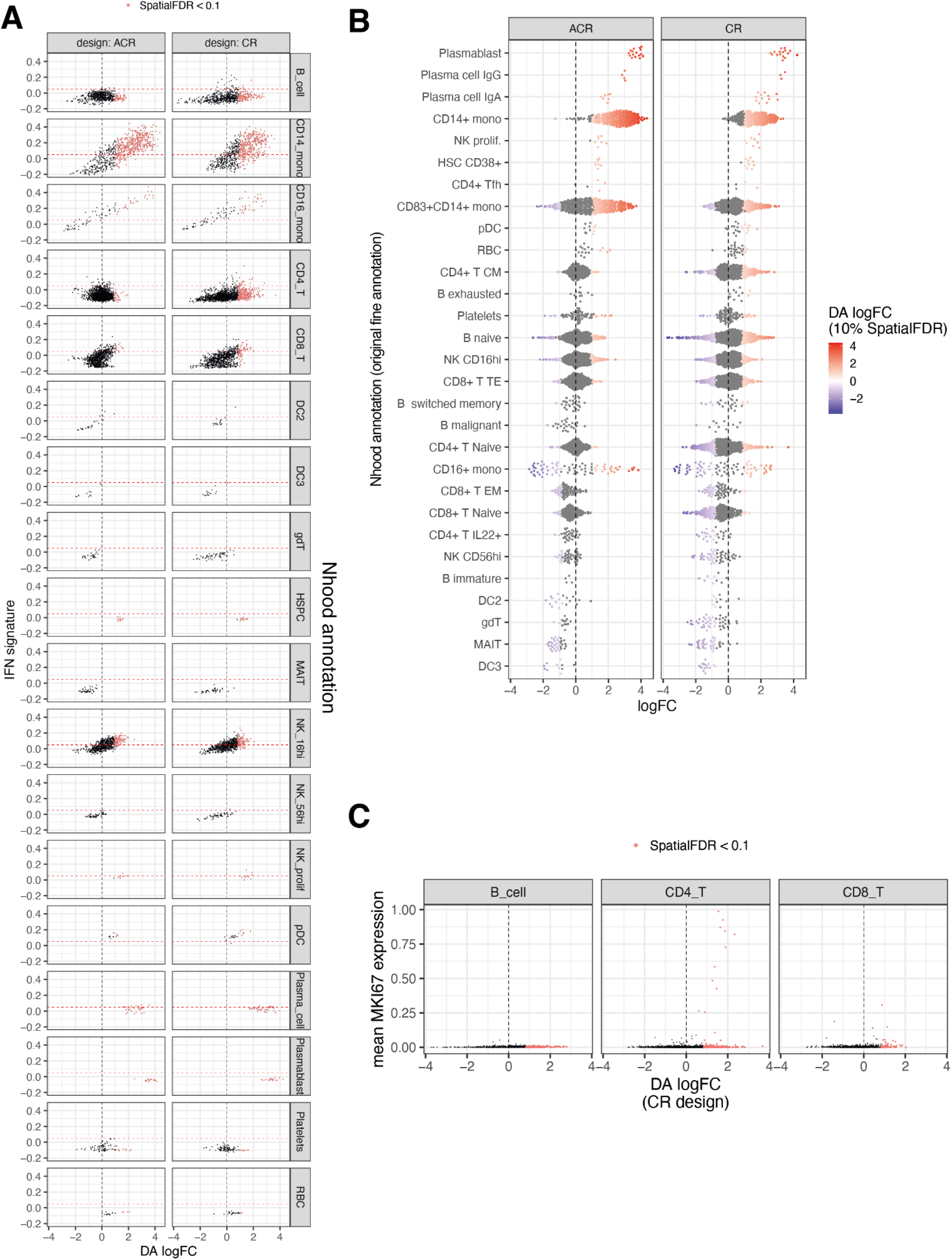
Reference design comparison on COVID-19 cohort. **(A)** Scatterplot of neighbourhood differential abundance log-Fold Change (DA logFC) against the mean expression of IFN signature with ACR design (left) and CR design (right), stratified by cell type annotation. Colored points indicate neighbourhoods where the enrichment was significant (10% SpatialFDR and logFC > 0). The dotted line denotes the threshold for high-IFN used for precision-recall analysis. **(B)** Beeswarm plot of DA logFC annotating neighbourhoods by fine annotation by Stephenson et al. Neighbourhoods where the differential abundance was significant (10% SpatialFDR) are colored. **(C)** Scatterplot of neighbourhood DA logFC against the mean expression of proliferation marker MKI67 in B and T cell states with CR design.

**Supplementary figure 9:**
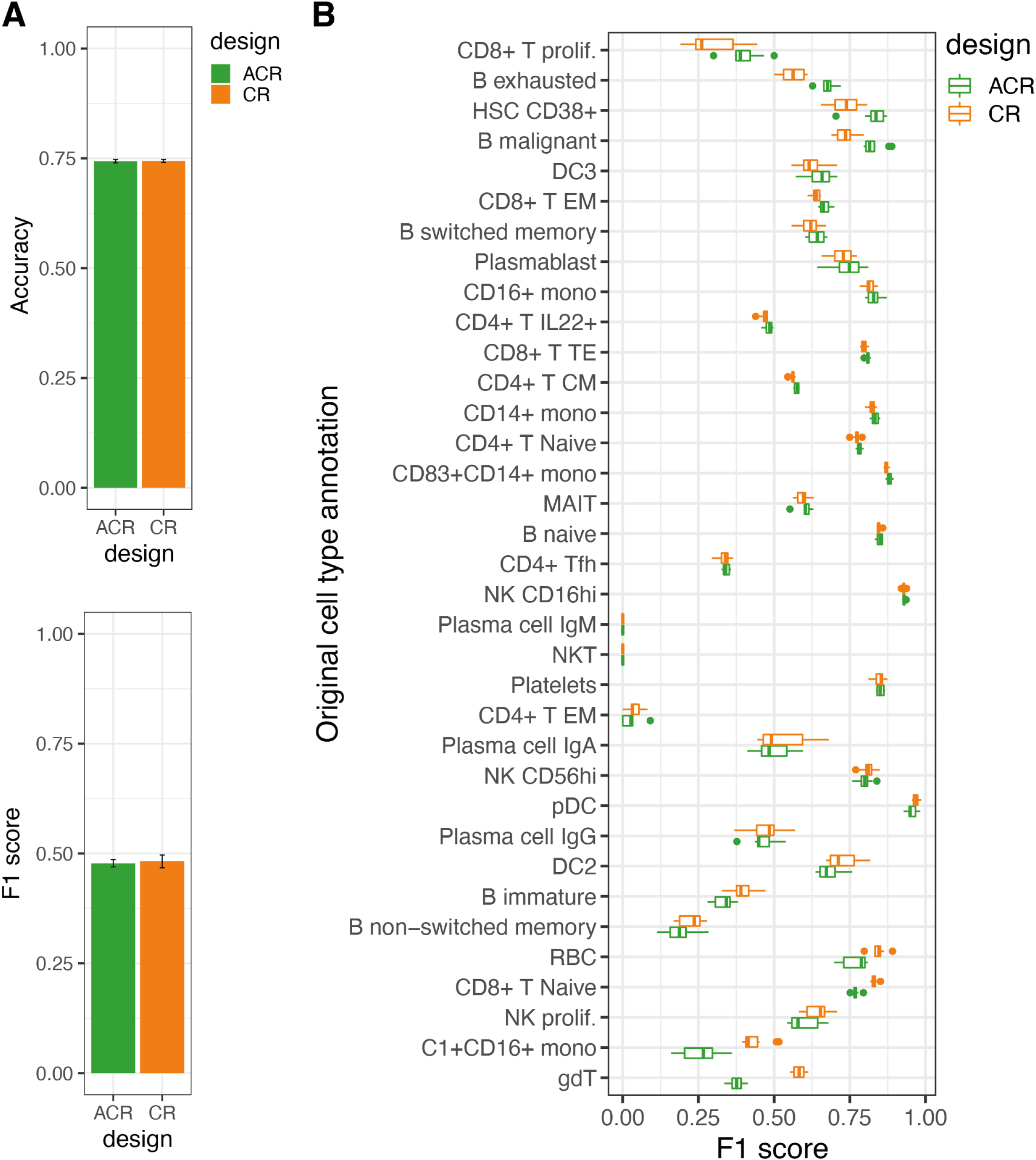
Latent space based classification of fine annotation labels. **(A)** Overall accuracy and mean F1 score for a KNN classifier for fine cell type labels from Stephenson et al. The classifier was trained on latent dimensions from ACR design and CR design. Cells from Stephenson et al. dataset (COVID-19 dataset and control dataset) were split at random in the training and test set (80% of cells in the training set). The mean performance on the test set over 10 different test-train splits is shown. The error bars show the standard deviation of the mean. **(B)** Cell type wise F1 score for KNN classifier trained on latent dimensions with different reference designs. Performance is shown only for cell types where at least 100 cells were found between the disease and control datasets. Cell types are ordered by the difference in median F1 score between designs.

**Supplementary figure 10:**
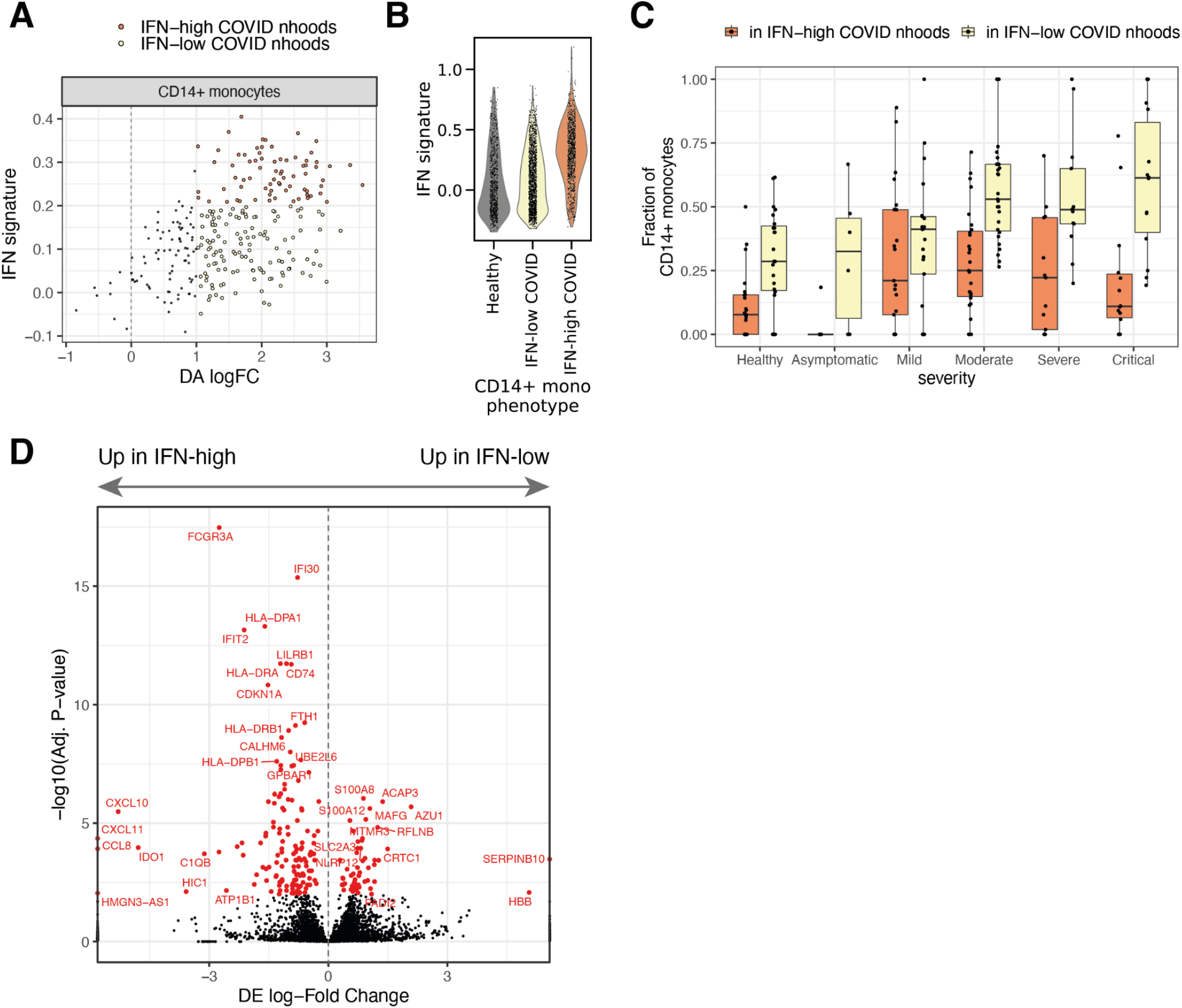
Heterogeneity in COVID-19 associated CD14+ monocyte states. **(A)** Scatterplot of neighbourhood differential abundance log-Fold Change (DA logFC) against the mean expression of IFN signature with CR design for neighbourhoods of CD14+ monocyte cells. Colored points indicate neighbourhoods where the enrichment in COVID-19 cells was significant (10% SpatialFDR and logFC > 0). Neighbourhoods are colored by IFN phenotype. **(B)** Distribution of IFN signature score for cells belonging to neighbourhoods in CR design assigned to 3 alternative CD14+ phenotypes. **(C)** Distribution of COVID-19 enriched CD14+ phenotypes (from CR design) across patients with varying disease severity: each point represents a donor, the y-axis shows the fraction of all CD14+ monocytes in that donor showing IFN-high COVID-19 enriched phenotype (orange), and IFN-low COVID-19 enriched phenotype (yellow). The remaining fraction are monocytes with healthy phenotype (not shown). **(D)** Volcano plot of differential expression analysis results from comparison between IFN-high and IFN-low COVID-19 specific CD14+ phenotypes identified with ACR design. Each point represents a tested gene, the x-axis shows the log-Fold Change of the differential expression test and the y-axis shows the adjusted p-value (using the Benjamini-Hochberg FDR correction). Genes where the difference was considered significant (FDR < 1%) are colored in red. A subset of significant genes are labelled.

## Supplementary Tables

**Suppl. Table 1: References of studies included in healthy PBMC dataset used for simulations**

**Suppl. Table 2: Differential expression analysis results for comparison of CD14+ monocytes COVID-19 phenotypes**

